# ALADIN is Required for the Production of Fertile Mouse Oocytes

**DOI:** 10.1101/043307

**Authors:** Sara Carvalhal, Michelle Stevense, Katrin Koehler, Ronald Naumann, Angela Huebner, Rolf Jessberger, Eric R. Griffis

## Abstract

Asymmetric cell divisions depend upon the precise placement of the mitotic spindle. In mammalian oocytes, spindles assemble close to the cell’s centre but chromosome segregation takes place at the cell periphery where half of the chromosomes are expelled into small, non-developing polar bodies at anaphases. By dividing so asymmetrically, most of the cytoplasmic content within the oocyte is preserved, which is critical for successful fertilization and early development. Recently, we determined that the nucleoporin ALADIN participates in spindle assembly in somatic cells, and we have also shown that female mice homozygous deficient for ALADIN are sterile. In this study we show that this protein is involved in specific meiotic stages including meiotic resumption, spindle assembly, and spindle positioning. In the absence of ALADIN, polar body extrusion is impaired in a majority of oocytes due to problems in spindle orientation prior to the first meiotic anaphase. Those few oocytes that can mature far enough to be fertilized in vitro are unable to support embryonic development beyond the two-cell stage. Overall, we find that ALADIN is critical for oocyte maturation and appears to be far more essential for this process than for somatic cell divisions.

## Introduction

Accurate chromosome segregation during mitotic and meiotic cell division is largely dependent on the proper assembly and orientation of a microtubule spindle (McNally, 2013; Ohkura, 2015). Spindle orientation determines where division takes place, and consequently the size of the daughter cells (McNally, 2013). In the large majority of animal cells, the spindle is located at the centre of the cell, and after division, two daughter cells of equal size are generated. Most asymmetric cell divisions are produced by displacing the spindle towards one side of the cell. Mouse oocytes have an extreme version of asymmetric spindle positioning (Brunet and Maro, 2007; Fabritius *et al*., 2011). During meiosis I, the spindle assembles in the middle of the cell, but then it migrates towards to the cortex. Once one of the spindle poles reaches the cortex, the spindle rotates and adopts a perpendicular orientation relative to the cortex (Fabritius *et al*., 2011; McNally, 2013). This event marks anaphase onset, where chromosomes are extruded into a small, non-developing polar body (Fabritius *et al*., 2011; McNally, 2013). This extremely asymmetric division is essential for fertilization and successful early embryonic divisions as it preserves almost all the cytoplasmic content within the oocyte (Fabritius *et al*., 2011; Chaigne *et al*., 2012).

In somatic cells, interactions between polar microtubules and the cortex regulate spindle positioning. Oocytes, however, lack centrosomes and polar microtubules to interact with the cortex, and the large size of oocytes relative to the spindle also makes it difficult for microtubules to interact with the cortex. Oocyte meiotic spindles therefore rely upon an alternative-positioning pathway based on the actin cytoskeleton (Verlhac *et al*., 2000; Dumont *et al*., 2007; Schuh and Ellenberg, 2008; Yi and Li, 2012), but the mechanisms that govern spindle assembly and positioning during oocyte maturation are still not fully understood (Brunet and Maro, 2007; Fabritius *et al*., 2011; Chaigne *et al*., 2012; Howe and FitzHarris, 2013; McNally, 2013).

Infertility affects at least 10% of couples, and between 20 to 30% of the cases are caused by failures in meiosis, which relies upon spindle assembly/positioning and chromosome alignment/segregation mechanisms that are very different from those that are utilized by somatic cells (Martin-du Pan and Campana, 1993; Lilford *et al*., 1994; Yeste *et al*., 2016). Thus, it is imperative to identify novel meiotic factors and to understand their roles, which was the motivation for this study. Mice homozygous for a deletion of the *Aaas* gene are viable; however, null females are sterile (Huebner *et al*., 2006), suggesting that its gene product ALADIN could be required for oocyte maturation and production of functional female gametes.

*Aaas* was first identified as the gene that is mutated in triple A syndrome, and later it was shown that the ALADIN protein is a component of the nuclear pore complex, which regulates nucleocytoplasmic transport (Handschug *et al*., 2001; Cronshaw *et al*., 2002). ALADIN mutations cause defects in the nuclear import of DNA repair proteins and ferritin during interphase (Storr *et al*., 2009; Prasad *et al*., 2013; Juhlen *et al*., 2015). Studies in human cells have now uncovered ALADIN’s role in spindle assembly via the spatial regulation of Aurora A kinase (Carvalhal *et al*., 2015). We showed that during mitosis ALADIN localizes around the spindle, and its depletion slows spindle assembly, weakens the spindle, and reduces spindle length (Carvalhal *et al*., 2015). Given the essential role of spindles in meiosis, we addressed a potential role of ALADIN in meiotic spindle formation and behaviour.

Here we show that ALADIN is essential for proper meiotic spindle assembly and positioning during meiosis in mouse oocytes, leading to catastrophic failures in the first meiotic divisions for most oocytes. We also show that the few ALADIN null oocytes that can be fertilized *in vitro* cannot support embryonic development beyond the two-cell stage, demonstrating that this protein is essential for producing fertile eggs.

## Results

### ALADIN’s localization is conserved during meiosis in mouse oocytes

During interphase, ALADIN is localized at the nuclear pore complex (Cronshaw *et al*., 2002). In mitotic human cells, ALADIN localizes around the mitotic spindle and enriches at spindle poles (Carvalhal *et al*., 2015). To test whether ALADIN’s localization is conserved during meiotic divisions, we fixed metaphase I and II wild-type mouse oocytes and stained them with an ALADIN specific antibody. During metaphase I, ALADIN mainly accumulated around the meiotic spindle (**Figure 1**A) and its localization was maintained throughout metaphase II (**Figure 1**A, lower two panels showing two representative images of oocytes at metaphase II). Additionally, when the spindle is near to the cortex, some ALADIN is still observed at the centre of the cell.

**Figure 1:**
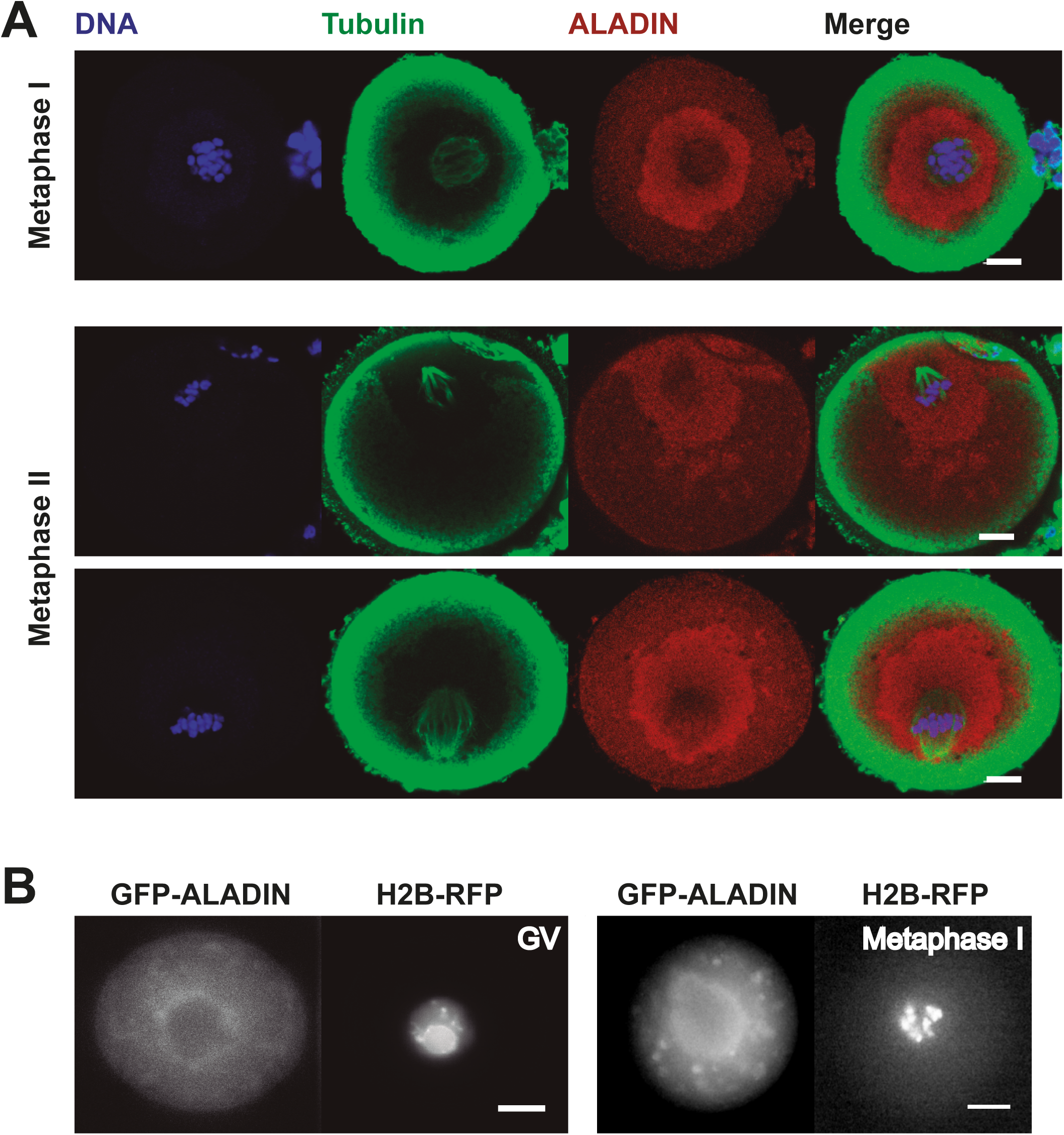
ALADIN localizes around the meiotic spindle during mouse oocyte maturation. (A) Wild-type and *Aaas*^-/-^ oocytes at metaphase I (one representative image) and II (two representative images) were isolated fixed and stained for tubulin, ALADIN and DNA. Representative Z-stack images. (B) Live imaging performed with wild-type mouse oocytes expressing GFP-ALADIN and H2B-RFP. Single Z-stack frames are presented from representative oocytes at germinal vesicle (GV) stage (left panel) and at metaphase I (right panel). Scale bars = 10 μm. ALADIN’s localization was observed in more than 3 independent experiments.

To confirm ALADIN’s localization, we injected WT oocytes with capped mRNAs encoding GFP-ALADIN and H2B-RFP and imaged developing oocytes up to metaphase I. Before nuclear envelope breakdown (NEBD), GFP-ALADIN was localized at the membrane of the germinal vesicle (GV; nucleus, **Figure 1**B). When chromosomes were fully condensed and aligned at the metaphase plate, ALADIN was localized around the aligned chromosomes in a distribution that was similar to the anti-ALADIN staining (**Figure 1**A). The presence of GFP-ALADIN was also observed in clusters throughout the cytoplasm, potentially a result of the overexpression of this protein. These results suggest a conservation of ALADIN’s localisation around the spindle in somatic and germ cells.

Interestingly, we found that the injection of a high concentration of ALADIN mRNA impairs GV breakdown (GVBD; Figure 2A). Injection of low levels of ALADIN mRNA into wild-type oocytes did not affect the timing of GVBD, but injection with twice the amount of mRNA resulted in 24 of the 49 oocytes failing to undergo GVBD 6 hours after meiosis resumption. However, the oocytes injected with the higher levels of mRNA that did undergo GVBD, did so with a timing that was similar to what was seen in control oocytes or oocytes injected with 45% less ALADIN mRNA. This binary response to the injection of high levels of ALADIN mRNA is consistent with an all or nothing defect in meiotic resumption rather than a specific defect in GVBD. This suggests that an overabundance of ALADIN can have a dominant negative effect in meiosis.

**Figure 2:**
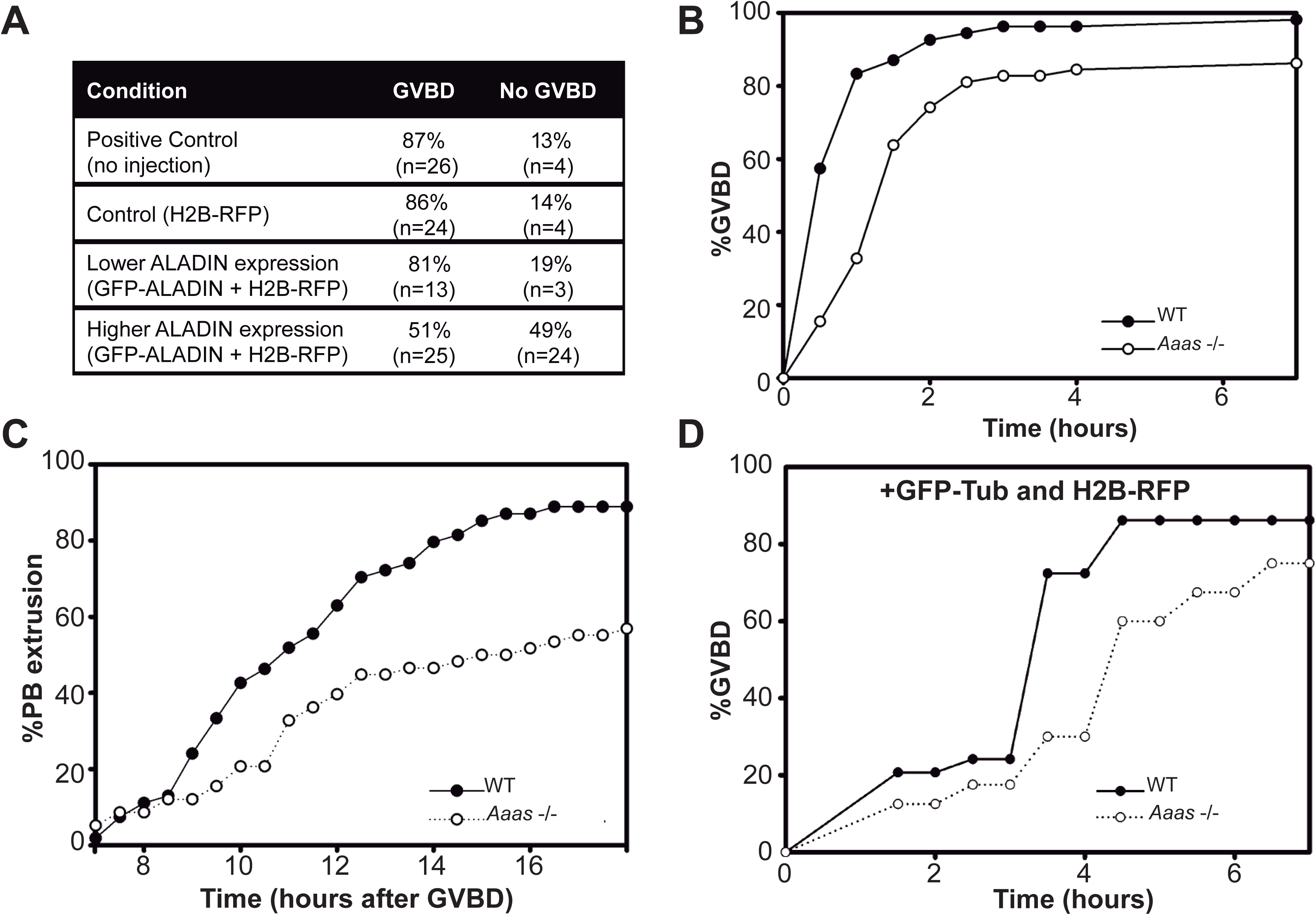
ALADIN levels are important for proper GVBD and polar body extrusion. (A) Wild-type oocytes were injected with the indicated mRNAs and the cumulative percentage of GVBD was measured over time. Uninjected oocytes were used as the positive control, and all the remaining conditions were injected with similar volumes of capped mRNAs. To express lower levels of ALADIN, oocytes were injected with a GFP-ALADIN mRNA 45% less concentrated (4.4 μg/μL) than in the higher expression (8 μg/μL) condition. The table shows the percentage of these oocytes that underwent GVBD. *n* indicates the number of oocytes measured in each condition. GVBD mean time ± standard deviation for Control: 2.2 ± 0.5 h; H2B-RFP: 2.4 ± 1.1 h; lower expression of GFP-ALADIN + H2B-RFP: 2.3 ± 0.4 h and higher expression of GFP-ALADIN + H2B-RFP: 2.62 ± 1.0 h. All experiments were performed at least 3 times with the exception of lower expression of GFP-ALADIN + H2B-RFP (n=1, result from oocytes collected from two mice). (B) Cumulative percentage of GVBD over time (h, hours) in wild-type (WT) and *Aaas*^-/-^ oocytes. More than 100 oocytes from 5 independent experiments were collected and measured for each condition evaluated. The average time for GVBD of WT oocytes was 1.9 ± 1.3 h while *Aaas*^-/-^ oocytes took significantly longer to achieve GVBD (2.7 ± 1.6 h, ***p<0.001). (C) Cumulative percentage of first polar body extrusion over time in WT and *Aaas*^-/-^ oocytes. n=5, more than 100 oocytes quantified. Mean ± standard deviation: 9.6 ± 2.9 h *vs* 11 ± 2.5 h; **p<0.007. (D) Cumulative percentage of GVBD over time (h, hours) in WT (control) and *Aaas*^-/-^ oocytes expressing GFP-β-Tubulin and H2B-RFP.

### Deletion of ALADIN disturbs oocyte maturation *in vitro*

Next we tested whether a lack of ALADIN could affect meiosis. ALADIN female homozygous knockout mice (*Aaas*^-/-^) were found to be viable but sterile. Histological sections of ovaries from wild-type and ALADIN-deficient mice did not show any differences, and follicular development and ovulation appeared to be normal (Huebner *et al*., 2006). Therefore, we decided to analyse if this infertility could be a consequence of ALADIN’s role in meiosis and the maturation process of oocytes. Wild-type (WT; *Aaas*^+/+^) and *Aaas*^-/-^ oocytes were collected and maintained in prophase arrest with intact GVs by incubation in milrinone containing buffer to arrest and synchronize them. Two hours after removal from milrinone and the onset of meiotic resumption, 91% of the 102 WT oocytes underwent germinal vesicle breakdown (GVBD), but only 58.5% of 105 *Aaas*^-/-^ oocytes had done so. On average, ALADIN-deficient oocytes required significantly longer to undergo GVBD (Figure 2B; 1.9 h ± 1.3 *vs* 2.7 h ± 1.6; ***p<0.001). 14 hours after GVBD, 80% of WT oocytes extruded their first polar body (PB). In contrast, only 46% of *Aaas*^-/-^ oocytes extruded a polar body, which increased to 57% 18 hours after GVBD (Figure 2C). This process was also significantly slower in the *Aaas*^-/-^ than the WT (Figure 2C; WT: 9.6 h ± 2.9 *vs *Aaas**^-/-^: 11 h ± 2.5; **p<0.007).

### ALADIN is required for correct spindle positioning in asymmetric oocyte divisions

So far it was shown that the absence of ALADIN slows both GVBD and PB extrusion timing and hinders the extrusion of PBs in oocytes. Our ALADIN depletion studies in somatic cells found that this protein is required for the timing of spindle assembly and formation of spindles of the proper length (Carvalhal *et al*., 2015). As ALADIN’s localization is conserved around the spindle in both cell models, we next tested if ALADIN could also participate in spindle assembly during oocyte cell division. Meiotic spindle assembly with and without ALADIN was analysed by time-lapse fluorescence imaging of tubulin/microtubules (GFP-β-Tubulin) and chromosomes (H2B-RFP) after injection with the respective capped mRNAs (Movie S1). As before, injected *Aaas*^-/-^ oocytes also took longer to achieve GVBD and had a reduced ability to extrude polar bodies (Figure 2D and S1A). These similarities suggested that neither increased tubulin expression nor the injection itself contributed to the *Aaas*^-/-^ phenotype. Representative frames of this time course are shown in Figure 3, which includes a representative example of a *Aaas*^-/-^ oocyte that successfully extruded its PB (middle column) and another that failed to form a PB (right column).

**Figure 3:**
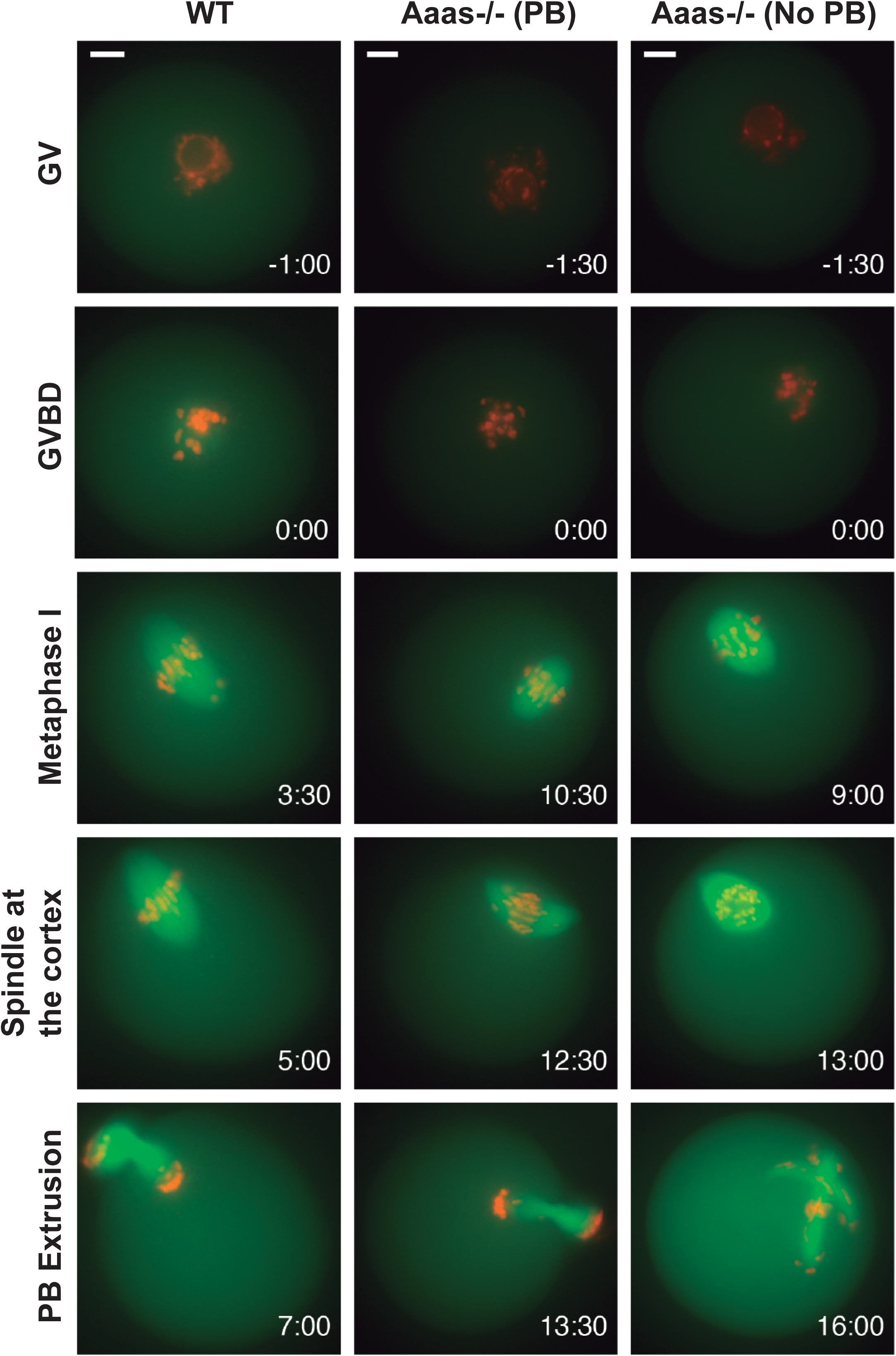
Oocytes lacking ALADIN show impaired spindle positioning and have less robust spindles. Time lapse analysis of progression through the first meiotic division in control (WT) and *Aaas*^-/-^ oocytes expressing GFP-β-Tubulin and H2B-RFP. Middle column shows a representative *Aaas*^-/-^ oocyte that was able to extrude a polar body. The right column shows representative *Aaas*^-/-^ oocyte that fails to extrude a polar body. Representative still frames presented. Time 0:00 represents GVBD in hours. GV = Germinal Vesicle; GVBD = Germinal Vesicle Breakdown and PB = Polar Body. Scale bars = 10 μm.

On average, *Aaas*^-/-^ oocytes took 96 minutes longer than wild-type oocytes to assemble a bipolar metaphase spindle after GVBD (Figure 4A; 3.0 h ± 1.9 vs 4.6 h ± 3.2; **p<0.05). Also, these bipolar structures were significantly shorter (Figure 4C; 8.3% shorter; **p<0.05) when compared with WT spindles.

**Figure 4:**
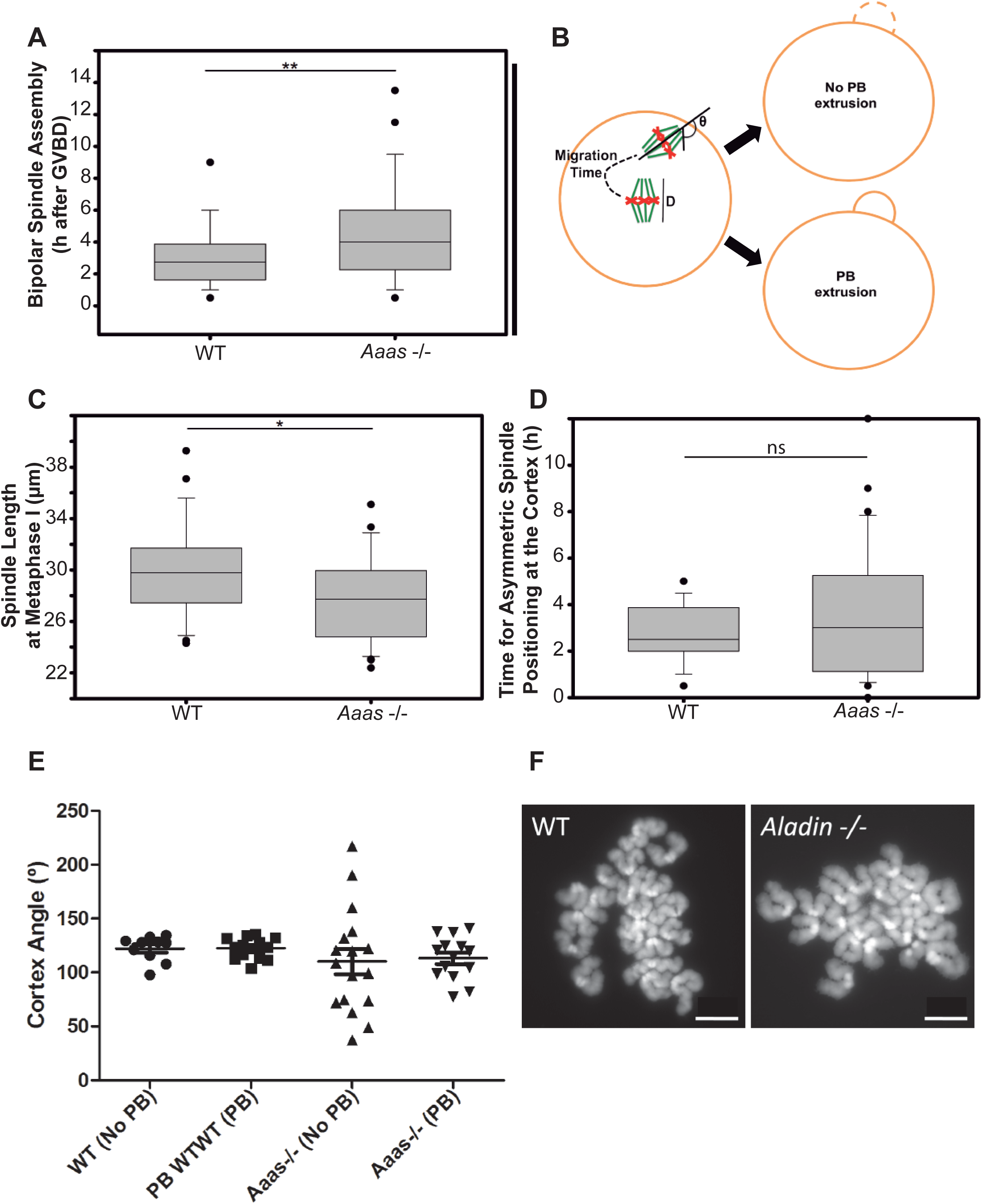
ALADIN is required for correct spindle assembly and positioning in oocyte meiosis. (A) The time from GVBD to bipolar spindle assembly was measured in control (WT) and *Aaas*^-/-^ oocytes expressing GFP-β-Tubulin and H2B-RFP. *Aaas*^-/-^ oocytes formed spindles more slowly and showed greater variability in spindle assembly timing (3.0 ± 1.9 h *vs* 4.6 ± 3.2 h; **p<0.005). Bipolar spindle assembly was scored when two individual poles were distinguishable. (B) Explanatory diagram of the measurements reported in C-E. The pole-pole distance, D, of metaphase I spindles were measured when bipolar spindles structures have a compact and organized metaphase plate. The black dotted line represents the migration time from the assembly of metaphase I spindles to the point when spindles contact the oocyte cortex. Θ represents the angle made between the spindle interacting with the cortex and a line that cross both spindle poles and the cortex. (C) Box-plot showing metaphase I spindle length in WT and *Aaas*^-/-^ oocytes expressing GFP-β-Tubulin and H2B-RFP. *Aaas*^-/-^ oocytes have a shorter spindle in metaphase; 29.9 ± 3.6 μm *vs* 27.5 ± 3.3 μm, **p<0.05. (D) The migration time of the spindle, as explained in (B), was measured for the indicated conditions (2.8 ± 1.2 h *vs* 3.5 ± 2.7 h). ns= no significant. (E) The cortex angle represented as Θ and explained in (B) was measured and the measurements are reported for oocytes that extruded a polar body (designated as PB) and oocytes that failed to extrude their polar body after reaching the cortex (No PB). WT PB: 122.3° ± 11.8 from 14 oocyte measured, WT NO PB: 122.5° ± 9.6 (10 oocytes), *Aaas*^-/-^ PB: 113.1° ± 19.9 (20 oocytes) and *Aaas*^-/-^ NO PB: 110.3° ± 48.4 (13 oocytes). All figures represent the results obtained from the analysis of four independent experiments executed with similar concentrations of GFP-β-Tubulin and H2B-RFP mRNAs (WT = 26 and *Aaas*^-/-^ = 36 oocytes analyzed). (F) Representative metaphase II chromosome spreads of WT and *Aaas*^-/-^ oocytes are presented. ALADIN deletion does not affect chromosome segregation after PB extrusion, since 20 sister chromatids are present in metaphase II. It also does not affect chromosome structure or sister chromatid pairing. Box-and-whisker plot: middle line shows the median value; the bottom and top of the box show the lower and upper quartiles (25-75%); whiskers extend to 10th and 90th percentiles, and all outliers are shown. **p<0.05 and ns = non-significant after t-test analysis. Scale bars = 10 μm.

Asymmetric positioning of meiotic spindles allows for the expulsion of chromosomes into small, non-developing polar bodies (Fabritius *et al*., 2011; McNally, 2013). To examine spindle relocation from the centre of the oocyte to the cortex, we measured the time from spindle bipolarisation to first contact with the cortex (Figure 4B and 4D). ALADIN null oocytes showed a slight slowing of this migration of the spindle to the cortex, when compared with WT oocytes (Figure 4D; 2.8h ± 1.2 vs 3.5 h ± 2.7).

Prior to anaphase, the spindles are attached by one pole to the oocyte cortex. To analyse how spindles approached the cortex, we measured the angle formed between the spindle pole and the cell periphery (Θ, Figure 4B and 4E) as it first interacted with the oocyte cortex. We sorted this data for cells that could extrude a polar body (PB) or for those that could not (NO PB). In wild-type conditions, independent of their success in extruding polar bodies (PB and NO PB), spindles consistently arrived at the oocyte cortex at a fairly defined angle (Figure 4E, 122.3° ± 11.8 and 122.5° ± 9.6, respectively and Movie S1). However, in the case of the *Aaas*^-/-^ oocytes we noticed differences in how the spindle approached the cortex (Movies S2 and S3). In *Aaas*^-/-^ ooctyes that ejected a polar body the spindles approached the cortex at an angle that was similar to what is seen in wild-type oocytes (113.1 ± 19.9 – 13 oocytes). In *Aaas*^-/-^ oocytes that failed polar body ejection, we observed a highly variable angle of approach between the spindle pole and the cortex (110.3° ± 48.4 – 20 oocytes). Movies S2 and S3 show examples of *Aaas*^-/-^ oocytes that ejected and did not eject a polar body respectively. In these oocytes we observe that the spindle can approach the cortex in an oblique, looping manner (in movie S2) but polar body ejection can proceed normally, while the spindle within the oocyte in movie S3 approaches the cortex in a very direct fashion but fails to anchor for polar body ejection. These results suggest that ALADIN is required for both stereotypical spindle migration as well as the formation of proper connections between the spindle and the oocyte cortex.

### ALADIN is essential for the production of fertile eggs

Our results suggest that ALADIN null oocytes could fail PB extrusion due to severe problems in the asymmetric positioning of the meiotic spindle relative to the oocyte cortex. However, some ALADIN depleted cells did successfully eject a PB, and so we tested if these oocytes were able to support fertilization and embryonic maturation.

We counted the chromosomes present in oocytes after PB ejection in chromosome spreads from wild-type and *Aaas*^-/-^ oocytes. Normal complements of chromosomes with grossly typical structures were found in both conditions (Figure 4F). These results reinforced the finding that some oocytes lacking ALADIN progress to a point where they can be fertilized.

To determine if these oocytes can be fertilized and support embryogenesis, we used laser-assisted and conventional *in vitro* fertilization (IVF; Figure S1). WT (Aaas^+/+^) spermatozoa were used to fertilize wild-type or *Aaas*^-/-^ matured oocytes. Fertilized embryos were then incubated until the first mitotic cleavage (two-cell embryo) becomes visible. In control conditions (WT matured oocytes + WT spermatozoa) several (15 of 27 in vitro matured oocytes fertilized by laser-assisted in vitro fertilization) formed two-cell stage embryos (representative embryos shown in Figure 5). However, in the absence of ALADIN (*Aaas*^-/-^ matured oocytes + WT spermatozoa), we only observed a single normal appearing 2-cell embryo out of the 15 laser-assisted in vitro fertilized oocytes (shown in the middle row; Figure 5). In the *Aaas*^-/-^ conditions, we also observed five embryos with a high degree of fragmentation and multinucleation. These morphological parameters are indicators of very poor quality embryos and could have resulted from the fertilization of oocytes that failed to eject chromosomes during an anaphase. To distinguish both embryo groups in *Aaas*^-/-^ conditions, the single normal appearing 2-cell embryo found was isolated from the others.

**Figure 5:**
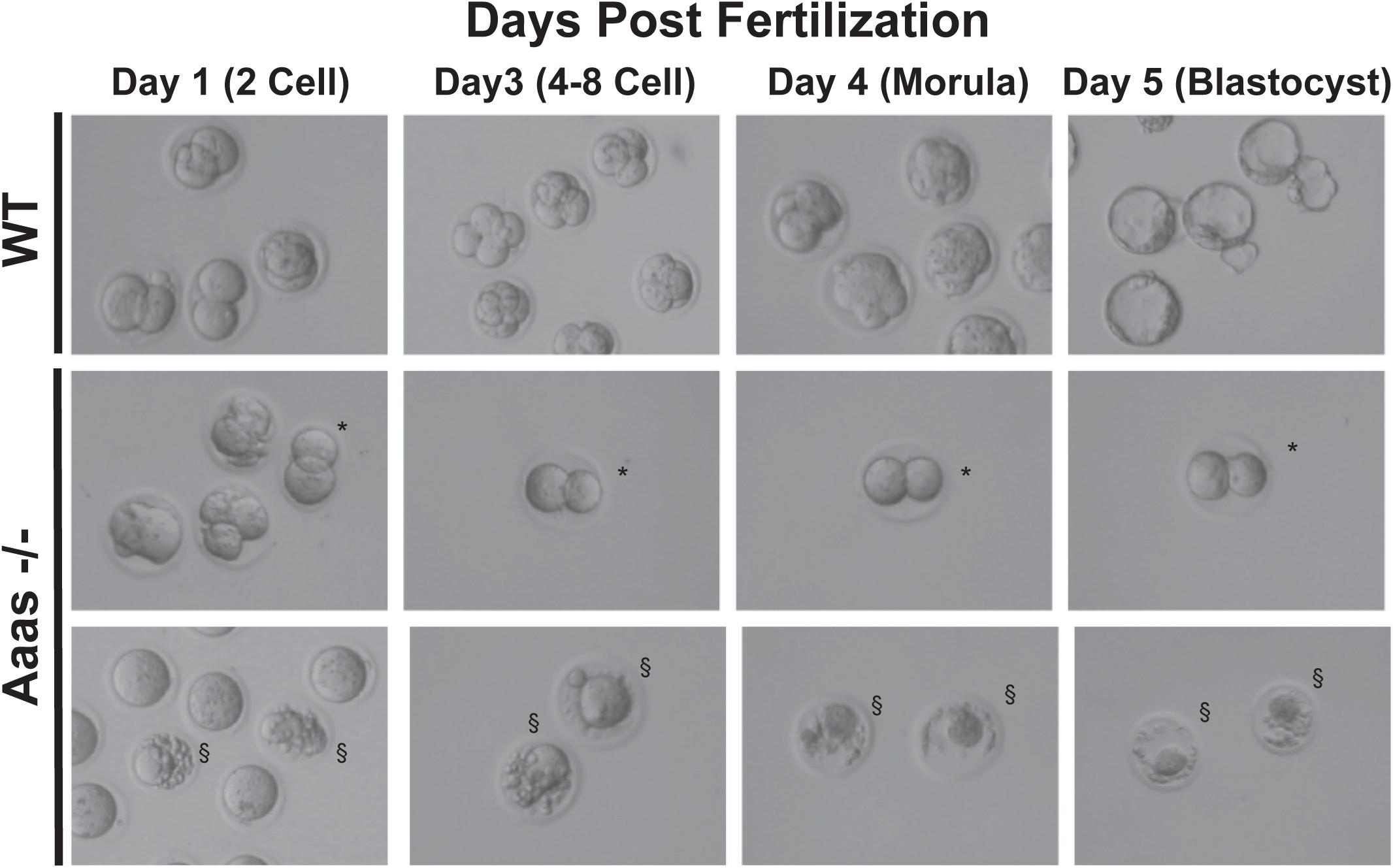
*Aaas*^-/-^ oocytes cannot support early embryogenesis. Laser-assisted and conventional IVF were used to fertilize WT and *Aaas*^-/-^ oocytes with WT spermatozoa. Two-cell embryos were isolated and embryogenesis was visualized for the next five days until the blastocyst stage. * indicates the single two-cell embryo observed in *Aaas*^-/-^ conditions (middle row). § shows representative abnormal morphologies, which were separated away from the two-cell embryo (bottom row).

The two-cell WT embryos all progressed to blastocysts and cavitated normally, confirming the success of IVF to form quality embryos. Conversely, the single *Aaas*^-/-^ equal 2-cell embryo did not progress further, and the abnormal *Aaas*^-/-^ embryos did not form blastocysts. These results suggest that although about 40% of *Aaas*^-/-^ oocytes are able to properly segregate their chromosomes in meiosis I and a portion of these can be fertilized, the resulting embryos cannot mature without ALADIN, explaining why these female mice are infertile.

## Discussion

Nucleoporins (NUPs) are components of the nuclear pore complex (NPC) that is essential for nuclear transport during interphase, but several nucleoporins are also known to be involved in chromosome segregation during mitosis (Chatel and Fahrenkrog, 2011; Mossaid and Fahrenkrog, 2015). Recently, we have shown that the NUP ALADIN has functions in mitotic spindle assembly in somatic cells from humans and fruit flies (Carvalhal *et al*., 2015). In line with functions during cell division, here we show that ALADIN also has roles in multiple steps of meiosis and is essential for producing oocytes that are capable of supporting embryogenesis.

After long periods in prophase I arrest, meiotic resumption and GVBD is triggered by the Maturation Promoting Factor (MPF) (Howe and FitzHarris, 2013). MPF enters the nucleus and phosphorylates NUPs promoting NPC disassembly (Margalit *et al*., 2005). Here we show that levels of ALADIN are important for GVBD timing and progression. An overabundance of ALADIN causes almost 50% of oocytes to arrest at the GV state. Although the involvement of ALADIN at NEBD in somatic cells has not yet been determined, ALADIN is tethered to the NPC by the transmembrane NUP NCD1 (Kind *et al*., 2009), which is required for NPC assembly in vertebrate cells (Haren *et al*., 2006). Therefore, we cannot exclude the possibility that ALADIN affects GVBD indirectly through altering NCD1 function or by altering the nuclear import of MPF. Alternatively, Aurora A has been shown to have a role in the initiation of meiosis in oocytes (Saskova *et al*., 2008); and we have shown that the overexpression of ALADIN can affect Aurora A’s activity in somatic cells (Carvalhal *et al*., 2015). Therefore, it is also possible that the overabundance of ALADIN could halt meiotic initiation and GVBD by reducing Aurora A activity. It was interesting to note that the overexpression of ALADIN had a very binary effect on GVBD. If cells did undergo GVBD after being injected with high levels of ALADIN mRNA, they did so with normal timing. This suggests that there may be a very sharp demarcation for the amount of ALADIN that will block GVBD and anything below this level does not affect this meiotic process.

ALADIN is present throughout the oocyte maturation process; it is localised at the NPC before GVBD, and when the spindle starts to assemble, ALADIN becomes more enriched around this structure. Therefore, GVBD likely marks a change of ALADIN’s function. So we cannot exclude that ALADIN’s role in GVBD timing is a cumulative effect of ALADIN’s sequential functions in interphase and mitosis.

Mouse oocytes lacking ALADIN show slower spindle formation, and the resulting spindles are shorter. Previously, we have shown similar defects on spindle assembly pathways in somatic cells (Carvalhal *et al*., 2015). These suggest that ALADIN’s functions are at least partially conserved between mitosis and meiosis. As in mitosis, we believe that ALADIN’s role is to promote robust spindle assembly. However, in the acentrosomal divisions of oocytes, we observe more severe defects than what we found in somatic cells.

In asymmetrically dividing somatic cells, astral microtubules generate pulling forces that displace the spindle towards one of the poles (McNally, 2013); in mouse oocytes spindle positioning and rotation is known to be driven by a cytoplasmic actin network (Dumont *et al*., 2007; Schuh and Ellenberg, 2008; Yi and Li, 2012). Our results indicate that spindle positioning and/or rotation is compromised in ALADIN-deficient oocytes. In *Aaas*^-/-^ oocytes, the migration of the spindle to the cell periphery is slower than in WT oocytes. Additionally, when the orientation of the spindle was compromised, these oocytes failed to eject a PB. No relationship between ALADIN and actin has been published so far, but ALADIN is localized around the spindle, where actin nucleation occurs and convection forces are generated (Grill *et al*., 2001; Bezanilla and Wadsworth, 2009).

A matrix or envelope has been observed to assemble around the spindle in many different systems. This structure is composed of nucleoporins, lamins and vesicles from the Golgi and ER and has been proposed to function as a non-microtubule scaffold able to concentrate spindle assembly factors, tether force generators and stabilise the mitotic spindle (Zheng, 2010; Jiang *et al*., 2014; Schweizer *et al*., 2014; Jiang *et al*., 2015a; Jiang *et al*., 2015b; Schweizer *et al*., 2015). Many nuclear pore proteins were identified by mass spectrometric analysis of purified *Xenopus* spindle matrices (Ma *et al*., 2009). When we consider ALADIN’s localization around the spindle and ability to interact with other NUPs, we think that it is reasonable to hypothesize that ALADIN could be a member of this matrix with a role in promoting spindle migration and rotation in the oocyte.

We found that only a small percentage of *Aaas*^-/-^ spindles reached the cortex with an approach angle that was similar to what we observed in control conditions. In these instances, oocytes were able to eject a polar body without problems in chromosome cohesion or structure, but we found that the quality of these mature oocytes is compromised, as they were unable to successfully develop into blastocyst embryos. At this point, we cannot exclude that ALADIN has additional roles in meiosis II or in the first mitotic division.

In conclusion, the female infertility seen in mice lacking ALADIN is caused by multiple defects in oocyte maturation, uncovering a new role of ALADIN in meiosis. Some oocytes can tolerate these perturbations and mature to the point where they can be fertilized, but the *Aaas*^-/-^ eggs are then incapable of supporting early embryogenesis. Mutations of ALADIN are known to cause triple A syndrome (Huebner *et al*., 2000; Sarathi and Shah, 2010). The majority of these patients develop their symptoms while very young, and fertility defects in these patients have never been closely analysed. Female mice that express only the S263P mutant variant of ALADIN are fertile (data not shown), suggesting that this mutant still retains some function in meiosis, which is consistent with our finding that cells from patients expressing only this variant have spindle assembly defects that are not as extreme as those seen in the Q387X variant expressing cells (Carvalhal *et al*., 2015). Interestingly, the *Aaas*^-/-^ mice show no defects in male fertility, suggesting that ALADIN plays a particularly important role in asymmetric and acentrosomal divisions. Our current data do not allow us to determine whether this is due to the fact that acentrosomal oocytes utilize an alternative spindle assembly and/or anaphase division pathway that relies more heavily on ALADIN or whether ALADIN is required for the proper preparation of the oocyte prior to the resumption of meiosis I. However, given that the acute depletion of ALADIN in somatic cells produces mechanical defects in spindle function and we also see defects in meiotic progression after ALADIN is overexpressed, we favor the former hypothesis.

## Materials and Methods

### Mouse oocyte and sperm collection and culture

C57BL/6 background WT (Aaas^+/+^) and *Aaas*^-/-^ were previously described (Huebner *et al*., 2006). Animals were bred and maintained under pathogen-free conditions at the Experimental Center of the Medizinisch-Theoretisches Zentrum of the Medical Faculty at the Dresden University of Technology according to approved animal welfare guidelines.

Oocytes were isolated from adult mice by puncturing the ovaries with needles in M2 medium supplemented with 2.5 μM milrinone to ensure prophase arrest. Surrounding follicle cells were removed by mouth pipetting. Resumption of oocyte maturation was induced by washing out the milrinone. For immunofluorescence analysis oocytes were incubated in M2 media covered by paraffin oil and maintained in a 5% CO_2_ atmosphere at 37°C with humidity control until the desired stage (metaphase I: 6-7 h, metaphase II: 12-13 h).

For *in vitro* fertilization (IVF) experiments, oocytes were also matured using superovulation techniques. Briefly, female mice (21-27 days of age) kept in a 12 hour light-dark cycle were intraperitoneal treated 44 h before the extraction (day 0) with 5 IU pregnant mare serum gonadotropin (PMSG) and 12 h before day 0 with 5 IU human chorionic gonadotropin (HCG). At day 0, oocytes were released from the cumulus complex. Spermatozoa were extracted from adult male mice by dissecting the epididymis and tail tip.

### *In vitro* transcription, microinjections, live oocyte imaging and quantitative analysis

GFP-β-Tubulin, Histone H2B-RFP, GFP-ALADIN mRNAs were synthesized with the T3 mMessage mMachine Kit (Ambion) according to the manufacturer's instructions, and purified using LiCl Precipitation (Ambion) or RNAeasy columns (Qiagen). The plasmids pRN3 GFP-Tubulin, Histone H2B-RFP were a gift from K. Wassmann (CNRS, Paris, France) (Touati *et al*., 2012). Human GFP-ALADIN was cloned into pRN3E (gift from K. Wassmann) with XhoI and BamHI to obtain GFP N-terminus-tagged ALADIN.

Oocytes arrested in prophase were injected using a FemtoJet microinjector (Eppendorf) with constant flow settings. To allow expression of fusion proteins and oocyte recovery, oocytes were incubated for 3-6 h in media supplemented with milirone. After release into M2 medium, oocytes were imaged every 30 min for up to 30 h on an inverted Nikon TE2000E microscope with a Plan APO 40×/1.25 NA objective or Plan APO 63×/1.4 NA objective, a CoolSNAP HQ camera (Photometrics), standard filter sets and with an environmental chamber to maintain 37°C. Z-series optical sections of 80 μm were recorded with slices taken every 7 μm For analysis of GVBD and polar body extrusion of non-injected oocytes a Plan APO 20×/0.75 NA objective was used. All images were acquired using the Nikon NIS-Elements ND2 software.

Quantitative analysis of time-lapse data was analysed manually using NIS-Elements software. Metaphase I time was defined when bipolar spindles structures have the most compact and organised metaphase plate. At this stage, pole-to-pole length of the spindle was measured using the tubulin signal. To calculate the migration of the spindle to the cortex, the time when one spindle pole reached the oocyte cortex was compared to the corresponding metaphase I timing. At this time, the angle made by the converging spindle pole with the cortex and spindle structure was measured.

### Immunofluorescence microscopy, quantitative analysis and antibodies used

Oocytes at the indicated maturation stage were briefly exposed to acid treatment (Tyrode's solution, Sigma) to partially remove the *zona pellucida* and excess acid was removed by prewarmed M2 medium. Then, oocytes were transferred to PHEM (60 mM PIPES, 25 mM HEPES, 10 mM EGTA and 2 mM MgCl_2_ pH 6.9) and incubated in 0.1% Triton X-100, 0.2% Glutaraldehyde, 2% PFA in PHEM 1x for 14 h at 4° C. After incubation, oocytes were transferred to Superfrost Plus Microscope Slides (Fisher Scientific) and allowed to attach at room temperature (25° C) without drying by adding excess PHEM. Oocytes were moved into a small region defined previously using nail polish to control the surface tension. After three 10 minute washes in PHEM-Wash (PHEM + 0.1% Triton X-100), oocytes were incubated in 1% Triton 1x PHEM for 30 min, followed by blocking in 3% BSA, 0.05% Tween 20 in 1x PHEM for 30 min at room temperature (25° C). Antibodies were diluted in the blocking solution and oocytes incubated with primary antibodies in a humidified chamber overnight at 4° C, and secondary antibody at room temperature for 30 min. Excess antibodies were removed after primary and secondary antibody incubations by three washes with PHEM–Wash for 10 min. Coverslips were mounted in Dako Fluorescence Mounting Medium (Dako) and sealed using nail polish.

Rat anti-α-tubulin (Pierce) was used at 1:200 and ALADIN (3E9, Santa cruz) at 1:50. Secondary antibodies were highly cross-subtracted Alexa Fluor-488, -555, conjugated antimouse, and ‐rat used at 1:200 (Life Technologies). 4',6-diamidino-2-phenylindole (DAPI) was used to stain chromosomal DNA at 2 μg/ml.

Fixed oocytes were imaged in a Leica SP2 confocal microscope and images processed in FIJI.

### Chromosome spreads

Chromosome spreads were prepared according to an adapted method from (Hodges and Hunt, 2002). Briefly, the zona pellucida of PB oocytes arrested at metaphase II was removed using acid treatment (Tyrodes solution; Sigma) and oocytes were transferred onto one well of a 10-well glass slide into 15μl of fixation solution (1% PFA, 0.1% Triton X-100, 3mM DTT; pH 9.2). After drying for several hours at room temperature, slides were washed 3 times 5 mins in PBS, and then mounted with Vectashield (Vector Laboratories) containing μg/ml DAPI (Sigma). Images were captured under oil using 100x magnification.

### Conventional and Laser-assisted *In Vitro* Fertilization (IVF)

For conventional IVF, ampulla from super ovulated mice were dissected from the oviduct and the cumulus-oocyte-complexes released, and incubated with fresh spermatozoa for 4 h at 37° C with humidity control and 5% CO_2_ atmosphere in Human Tubal Fluid medium. After incubation, oocytes were washed into KSOM mouse embryo culture medium to remove spermatozoa and cumulus cells and the fertilisation status of each oocyte was assessed by the presence of pronuclei 10 h after the beginning of the insemination. All oocytes were washed and transferred to pre-equilibrated KSOM medium and incubated at 37° C. Concentration and motility of fresh spermatozoa was evaluated and only spermatozoa with standard qualities were used. After overnight culture of the oocytes, 2-cell stage embryos were transfer into KSOM and their development monitored.

For laser assisted IVF, oocytes were harvested from ovaries, matured *in vitro*, and perforated with a 600 μs pulse of a 1460 nm laser set at 63% power at room temperature (Hamilton Thorne). Perforations took approximately 10 mins per conditions. Perforated oocytes were washed and transferred into an IVF drop containing spermatozoa, as described above.

### Statistical analysis and data representation

Data obtained from NIS-Elements software was imported into Sigma-plot software (Systat Software Inc), which was used to generate box-and-whisker and cumulative distribution plots, with the exception of the one showed in Figure 4E (GraphPad Prism). In box-and-whisker plots the middle line shows the median value; the bottom and top of the box show the lower and upper quartiles; whiskers extend to 10th and 90th percentiles, and all outliers are shown. Student’s t-test was used to determine statistical significance between two different treatments (significance is reported in each figure). Images represented were produced in Illustrator, FIJI and OMERO WebFigure.

## Acknowledgments

We thank Dr Kajta Wassmann for plasmids and Prof Attila Toth for discussion. We thank the Light Microscopy Facility, College of Life Sciences, University of Dundee, for help with imaging. MS is supported through institutional funds and DFG grant STE 2280/2-1, and AH and KK are supported by DFG grants HU 895/5-1 and HU 895/5-2. SC was supported by an EMBO Short Term Fellowship (ASTF 521-2014) and a Biotechnology and Biological Sciences Research Council studentship. RJ is supported by institutional funds and by DFG grant JE150/11-2. ERG was supported by a Wellcome Trust RCDF award (090064/Z/09/Z) and a Wellcome Trust Strategic award to the Centre for Gene Regulation and Expression (097945/B/11/Z).

## Author contributions

SC, ERG, and RJ designed the experiments. SC performed the majority of experiments, analysed the data and co-wrote the manuscript with ERG. MS performed chromosome spreads and helped in the design of experiments, microinjections and live imaging. KK performed all mouse genotyping and helped with fixed imaging and IVF. RN helped in IVF. AH and RJ contributed tools and reagents. MS, KK, AH and RJ contributed towards the manuscript.

---

**Figure S1:**
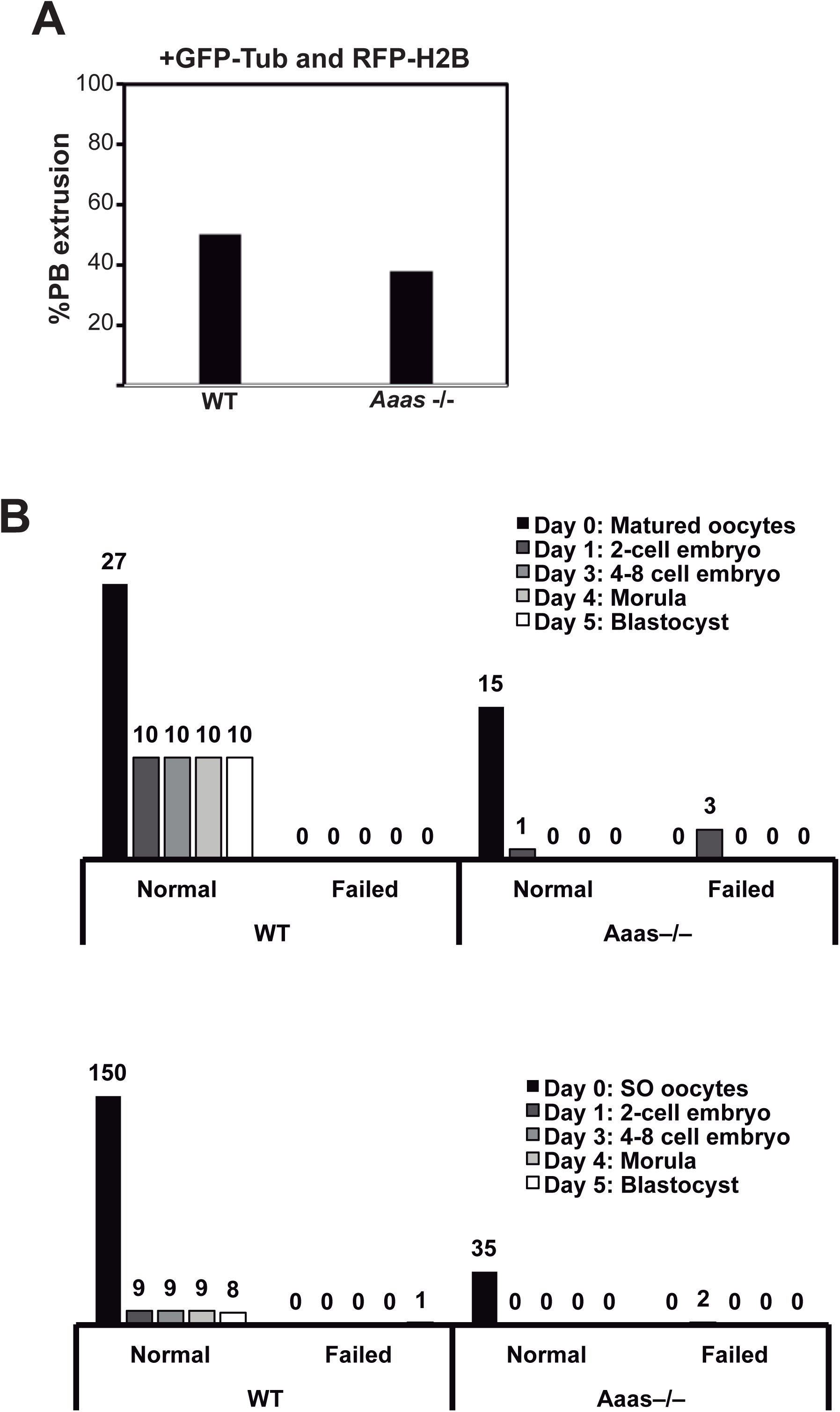
(A) Polar body extrusion was quantitated for WT and *Aaas*^-/-^ oocytes injected with mRNAs to induce the expression of GFP-β-Tubulin and H2B-RFP. Confocal Z-stacks of the oocytes were taken every thirty minutes; 50% of WT oocytes and 38.0% of *Aaas*^*-/-*^ oocytes ejected a polar body. Data obtained from four independent experiments executed with similar concentrations of GFP-β-Tubulin and H2B-RFP mRNAs (WT = 26 and *Aaas*^-/-^ = 36 oocytes). (B) Quantification of embryo stages during development of WT and *Aaas*^-/-^ oocytes fertilized with WT spermatozoa. Top panel: Laser-assisted IVF - three adult female mice of the indicated genotypes were induced to superovulate and their oocytes were collected after sacrifice. Only oocytes with visible polar bodies were used for IVF. Bottom panel: Conventional IVF - superovulation (SO) was induced in WT and *Aaas*^-/-^ female mice. None of the *Aaas*^-/-^ eggs could successfully be fertilized and generate blastocysts. Absolute numbers are presented for each stage.

**Movie S1:** Time-lapse analysis shows the progression through the first meiotic division in WT oocytes expressing GFP-β-Tubulin and H2B-RFP. The movie starts just before germinal vesicle breakdown occurs. In prometaphase I, the spindle is assembled and most chromosomes are aligned in the center of the oocyte. Then the spindle migrates to the cell cortex where it expels half its chromosomal contents into a polar body. Image stacks were acquired every 30 minutes and a maximum intensity projection is shown for each time point. Movie is shown at 9000x real time.

**Movie S2:** Successful polar body ejection in a *Aaas*^-/-^ oocyte. Z-stacks were collected every 30 minutes of *Aaas*^-/-^ oocytes expressing GFP-β-tubulin and H2B-RFP. Although this oocyte successfully ejects a polar body, the migration of the spindle to the cortex is not as direct as what is seen in the WT case in Movie S1. Z-stacks were converted to maximum intensity projections and are shown at 9000x real time.

**Movie S3:** Polar body ejection failure in a *Aaas*^-/-^ oocyte. Z-stacks were collected every 30 minutes of *Aaas*^-/-^ oocytes expressing GFP-β-tubulin and H2B-RFP. Z-stacks were converted to maximum intensity projections and are shown at 9000x real time.

## References

Bezanilla, M., and Wadsworth, P. (2009). Spindle positioning: actin mediates pushing and pulling. Current biology: CB 19, R168-169.

Brunet, S., and Maro, B. (2007). Germinal vesicle position and meiotic maturation in mouse oocyte. Reproduction 133, 1069-1072.

Carvalhal, S., Ribeiro, S.A., Arocena, M., Kasciukovic, T., Temme, A., Koehler, K., Huebner, A., and Griffis, E.R. (2015). The Nucleoporin ALADIN Regulates Aurora A Localization to Ensure Robust Mitotic Spindle Formation. Molecular biology of the cell.

Chaigne, A., Verlhac, M.H., Terret, M.E. (2012). Spindle positioning in mammalian oocytes. Experimental cell research 318, 1442-1447.

Chatel, G., and Fahrenkrog, B. (2011). Nucleoporins: leaving the nuclear pore complex for a successful mitosis. Cell Signal 23, 1555-1562.

Cronshaw, J.M., Krutchinsky, A.N., Zhang, W., Chait, B.T., and Matunis, M.J. (2002). Proteomic analysis of the mammalian nuclear pore complex. The Journal of cell biology 158, 915-927.

Dumont, J., Million, K., Sunderland, K., Rassinier, P., Lim, H., Leader, B., and Verlhac, M.H. (2007). Formin-2 is required for spindle migration and for the late steps of cytokinesis in mouse oocytes. Developmental biology 301, 254-265.

Fabritius, A.S., Ellefson, M.L., and McNally, F.J. (2011). Nuclear and spindle positioning during oocyte meiosis. Current opinion in cell biology 23, 78-84.

Grill, S.W., Gonczy, P., Stelzer, E.H., and Hyman, A.A. (2001). Polarity controls forces governing asymmetric spindle positioning in the Caenorhabditis elegans embryo. Nature 409, 630-633.

Handschug, K., Sperling, S., Yoon, S.J., Hennig, S., Clark, A.J., and Huebner, A. (2001). Triple A syndrome is caused by mutations in AAAS, a new WD-repeat protein gene. Hum Mol Genet 10, 283-290.

Haren, L., Remy, M.H., Bazin, I., Callebaut, I., Wright, M., and Merdes, A. (2006). NEDD1-dependent recruitment of the gamma-tubulin ring complex to the centrosome is necessary for centriole duplication and spindle assembly. The Journal of cell biology 172, 505-515.

Hodges, C.A., and Hunt, P.A. (2002). Simultaneous analysis of chromosomes and chromosome-associated proteins in mammalian oocytes and embryos. Chromosoma 111, 165-169.

Howe, K., and FitzHarris, G. (2013). Recent insights into spindle function in mammalian oocytes and early embryos. Biology of reproduction 89, 71.

Huebner, A., Mann, P., Rohde, E., Kaindl, A.M., Witt, M., Verkade, P., Jakubiczka, S., Menschikowski, M., Stoltenburg-Didinger, G.., and Koehler, K. (2006). Mice lacking the nuclear pore complex protein ALADIN show female infertility but fail to develop a phenotype resembling human triple A syndrome. Molecular and cellular biology 26, 1879-1887.

Huebner, A., Yoon, S.J., Ozkinay, F., Hilscher, C., Lee, H., Clark, A.J., and Handschug, K. (2000). Triple A syndrome‐‐clinical aspects and molecular genetics. Endocrine research 26, 751-759.

Jiang, H., He, X., Feng, D., Zhu, X., and Zheng, Y. (2015a). RanGTP aids anaphase entry through Ubr5-mediated protein turnover. The Journal of cell biology 211, 7-18.

Jiang, H., He, X., Wang, S., Jia, J., Wan, Y., Wang, Y., Zeng, R., Yates, J., 3rd, Zhu, X., and Zheng, Y. (2014). A microtubule-associated zinc finger protein, BuGZ, regulates mitotic chromosome alignment by ensuring Bub3 stability and kinetochore targeting. Developmental cell 28, 268-281.

Jiang, H., Wang, S., Huang, Y., He, X., Cui, H., Zhu, X., and Zheng, Y. (2015b). Phase Transition of Spindle-Associated Protein Regulate Spindle Apparatus Assembly. Cell 163, 108-122.

Juhlen, R., Idkowiak, J., Taylor, A.E., Kind, B., Arlt, W., Huebner, A., and Koehler, K. (2015). Role of ALADIN in human adrenocortical cells for oxidative stress response and steroidogenesis. PLoS One 10, e0124582.

Kind, B., Koehler, K., Lorenz, M., and Huebner, A. (2009). The nuclear pore complex protein ALADIN is anchored via NDC1 but not via POM121 and GP210 in the nuclear envelope. Biochemical and biophysical research communications 390, 205-210.

Lilford, R., Jones, A.M., Bishop, D.T., Thornton, J., and Mueller, R. (1994). Case-control study of whether subfertility in men is familial. Bmj 309, 570-573.

Ma, L., Tsai, M.Y., Wang, S., Lu, B., Chen, R., Iii, J.R., Zhu, X., and Zheng, Y. (2009). Requirement for Nudel and dynein for assembly of the lamin B spindle matrix. Nature cell biology 11, 247-256.

Margalit, A., Vlcek, S., Gruenbaum, Y., and Foisner, R. (2005). Breaking and making of the nuclear envelope. Journal of cellular biochemistry 95, 454-465.

Martin-du Pan, R.C., and Campana, A. (1993). Physiopathology of spermatogenic arrest. Fertility and sterility 60, 937-946.

McNally, F.J. (2013). Mechanisms of spindle positioning. The Journal of cell biology 200, 131-140.

Mossaid, I., and Fahrenkrog, B. (2015). Complex Commingling: Nucleoporins and the Spindle Assembly Checkpoint. Cells 4, 706-725.

Ohkura, H. (2015). Meiosis: an overview of key differences from mitosis. Cold Spring Harbor perspectives in biology 7.

Prasad, R., Metherell, L.A., Clark, A.J., and Storr, H.L. (2013). Deficiency of ALADIN impairs redox homeostasis in human adrenal cells and inhibits steroidogenesis. Endocrinology 154, 3209-3218.

Sarathi, V., and Shah, N.S. (2010). Triple-A syndrome. Advances in experimental medicine and biology 685, 1-8.

Saskova, A., Solc, P., Baran, V., Kubelka, M., Schultz, R.M., and Motlik, J. (2008). Aurora kinase A controls meiosis I progression in mouse oocytes. Cell cycle 7, 2368-2376.

Schuh, M., and Ellenberg, J. (2008). A new model for asymmetric spindle positioning in mouse oocytes. Current biology: CB 18, 1986-1992.

Schweizer, N., Pawar, N., Weiss, M., and Maiato, H. (2015). An organelle-exclusion envelope assists mitosis and underlies distinct molecular crowding in the spindle region. The Journal of cell biology 210, 695-704.

Schweizer, N., Weiss, M., and Maiato, H. (2014). The dynamic spindle matrix. Current opinion in cell biology 28, 1-7.

Storr, H.L., Kind, B., Parfitt, D.A., Chapple, J.P., Lorenz, M., Koehler, K., Huebner, A., and Clark, A.J. (2009). Deficiency of ferritin heavy-chain nuclear import in triple a syndrome implies nuclear oxidative damage as the primary disease mechanism. Molecular endocrinology 23, 2086-2094.

Touati, S.A., Cladiere, D., Lister, L.M., Leontiou, I., Chambon, J.P., Rattani, A., Bottger, F., Stemmann, O., Nasmyth, K., Herbert, M., and Wassmann, K. (2012). Cyclin A2 is required for sister chromatid segregation, but not separase control, in mouse oocyte meiosis. Cell Rep 2, 1077-1087.

Verlhac, M.H., Lefebvre, C., Guillaud, P., Rassinier, P., and Maro, B. (2000). Asymmetric division in mouse oocytes: with or without Mos. Current biology: CB 10, 1303-1306.

Yeste, M., Jones, C., Amdani, S.N., Patel, S., and Coward, K. (2016). Oocyte activation deficiency: a role for an oocyte contribution? Human reproduction update 22, 23-47.

Yi, K., and Li, R. (2012). Actin cytoskeleton in cell polarity and asymmetric division during mouse oocyte maturation. Cytoskeleton 69, 727-737.

Zheng, Y. (2010). A membranous spindle matrix orchestrates cell division. Nature reviews. Molecular cell biology 11, 529-535.

